# A Multilayer Human Skin Diseasome Links Molecular Architecture, Clinical Co-occurrence, and Drug-Reversal Signatures

**DOI:** 10.64898/2026.10.07.756451

**Authors:** Rashid Karim, Debajyoti Ghosh

**Author notes:** Correspondence: **D. Ghosh MSc., Ph.D.** Assistant Professor (Research), Division of Rheumatology, Allergy & Immunology. University Of Cincinnati College of Medicine Medical Science Building, Room 7410, 231 Albert Sabin Way Cincinnati, OH 45267.

## Abstract

**Background:** Immune-mediated skin diseases frequently co-occur, but the molecular architecture connecting clinically distinct disorders, and the extent to which molecular proximity predicts shared therapeutic opportunity, remain incompletely defined. We constructed a multilayer skin diseasome integrating transcriptomic overlap, pathway enrichment, protein-interactome topology, curated disease associations, real-world clinical co-occurrence, and drug-perturbation signatures across psoriasis (PS), atopic dermatitis (AD), hidradenitis suppurativa (HS), vitiligo (VL), and chronic spontaneous urticaria (CSU), using sodium lauryl sulfate-induced irritant contact dermatitis (SLS-ICD) as a mechanistically distinct barrier-injury comparator.

**Methods:** Differentially expressed genes (DEGs) were derived from publicly available NCBI Gene Expression Omnibus datasets and compared using exclusive pairwise and higher-order intersections. Gene Ontology biological-process enrichment characterized shared and condition-specific programs. Interactome-based network separation (S_AB) quantified the relative topological proximity of disease modules. DisGeNET enrichment generated candidate disease +associations, a selected subset of which was examined descriptively for patient co-occurrence in TriNetX. Candidate compounds were ranked by reversal of disease-associated expression signatures against LINCS L1000 perturbation profiles.

**Results:** The largest exclusive DEG intersection occurred between PS and HS (2,060 genes), followed by AD-PS-HS (490), VL-PS (464), AD-PS (402), and VL-PS-HS (379). Fourteen genes were shared across PS, AD, HS, VL, and CSU but not SLS-ICD. Neutrophil-associated processes recurred across the immune-mediated conditions, whereas SLS-ICD was dominated by keratinization and epidermal-differentiation programs. PS and HS had the lowest observed network-separation value (S_AB = −0.594); SLS-ICD had positive S_AB values in all five comparisons, although this separation did not persist once module sizes were harmonized. DisGeNET linked the index conditions to diverse immune, cardiovascular, neurologic, and neoplastic categories; 42 of these candidate diseases were observed to co-occur with an index condition in the queried TriNetX cohorts. Drug-signature overlap was greatest between PS and CSU, despite their marginal interactome separation, indicating that interactome proximity and predicted pharmacologic reversal capture different dimensions of disease relatedness. mTOR/PI3K-pathway inhibitors were prominent among the psoriasis reversal candidates, while methotrexate and mycophenolate mofetil emerged from both the CSU signature and the curated gene set associated with HIV infection.

**Conclusion:** Multilayer integration identified a connected molecular core among five immune-mediated skin diseases and a distinct SLS-ICD injury module. The discordance between interactome proximity and drug-signature overlap suggests that no single layer adequately defines disease relatedness. These findings provide a hypothesis-generating framework for prioritizing disease links and repurposing candidates, but controlled epidemiologic analyses and experimental validation are required before clinical or therapeutic inference.

## 1. Introduction

Human diseases are conventionally classified as discrete entities, yet the molecular programs that generate disease phenotypes often overlap. Network medicine represents diseases as perturbations of interconnected genes, pathways, and protein interactions, providing a framework for explaining reproducible comorbidity and for identifying shared therapeutic vulnerabilities. A central unresolved question, however, is whether different measures of disease relatedness such as transcriptomic overlap, pathway convergence, interactome proximity, clinical co-occurrence, and drug-signature reversal, can identify the same relationships or capture distinct dimensions of disease biology.

Clinically distinct diseases can share differentially regulated genes, pathways, and protein-interaction neighborhoods, forming interconnected disease networks that may help explain patterns of comorbidity. However, whether different measures of disease relatedness (e.g., transcriptomic overlap, pathway convergence, interactome proximity, clinical co-occurrence, and drug-signature similarity) identify the same relationships remains unclear. Discordance among these measures may reveal disease connections and therapeutic opportunities that would be missed by any single approach.

Skin diseases rarely occur in isolation. Psoriasis is associated with psoriatic arthritis, Crohn disease, and cardiometabolic disorders, while atopic dermatitis can precede asthma and allergic rhinitis in the atopic march.^1,2^ Epidemiologic associations have also been reported between PS and HS, HS and AD, AD and VL, and CSU and both PS and VL, suggesting that immune-mediated skin diseases may form a partially interconnected clinical network. In contrast, sodium lauryl sulfate-induced irritant contact dermatitis (SLS-ICD) is an experimental model of chemically induced barrier injury rather than a clinical disease with an established comorbidity spectrum. We therefore included SLS-ICD as a mechanistically distinct comparator, enabling us to ask whether the integrated framework can distinguish shared immune-disease architecture from acute irritant injury.

Network medicine interprets disease as a perturbation of an interconnected cellular system rather than the consequence of a single defective gene.^3^ Goh et al. constructed a human disease network by linking disorders that share associated genes and showed that genes associated with phenotypically similar disorders are more likely to interact physically and exhibit correlated expression.^4^ Gene-sharing approaches can nevertheless miss disorders whose molecular modules are topologically close without containing the same annotated genes. Menche et al. addressed this limitation with the network-separation statistic S_AB, which compares the mean shortest-path distance between two disease modules with the internal distances of each module.^5^ Negative values indicate relative topological overlap, whereas positive values indicate separation within the reference interactome.

Here, we used Menche et al.’s S_AB statistic as one layer of molecular evidence. S_AB quantifies relative topological proximity within a protein–protein interaction network; it does not establish causal propagation, biological equivalence, or clinical comorbidity. We therefore interpreted S_AB together with transcriptomic, pathway, clinical co-occurrence, and drug-perturbation evidence rather than as a stand-alone measure.

Network approaches have supported drug-repurposing studies in cardiovascular and other diseases, including analyses evaluated against large patient populations and perturbational screens across multiple disorders. ^6,7^ Yet common inflammatory and immune-mediated skin diseases have not been compared within a single framework that explicitly tests whether distinct molecular and translational layers converge. Individual diseases have well-described transcriptomic and comorbidity profiles, but their relative positions across gene-expression, pathway, interactome, clinical, and pharmacologic spaces remain incompletely mapped.

We therefore constructed a multilayer diseasome for PS, AD, HS, VL, and CSU, with SLS-ICD serving as a barrier-injury comparator. The analysis proceeded from DEG overlap and pathway enrichment to interactome separation, DisGeNET disease-association mapping, descriptive TriNetX co-occurrence, and LINCS L1000 drug-signature reversal. Our prespecified conceptual question was not only which conditions appeared molecularly close, but also whether proximity was preserved across layers and translated into similar candidate therapeutic signatures.

Interestingly, these six diseases collectively span a spectrum of etiologic architectures: from irritant contact dermatitis, in which environmental/chemical exposure predominates, through hidradenitis suppurativa, atopic dermatitis, psoriasis, and vitiligo, in which twin studies indicate substantial heritability (∼66–90%), to chronic spontaneous urticaria, in which genetic and environmental factors contribute more evenly (∼45–60% heritability).^8–12^ In particular, vitiligo shows 23% concordance rate in monozygotic twins and 0% in dizygotic twins indicating a multifactorial role. Collectively, these conditions span clinically and mechanistically diverse phenotypes. Rather than assuming that etiologic similarity would determine network proximity, we used their diversity to test whether independent molecular layers captured a coherent immune-mediated core and distinguished it from experimentally induced barrier injury. This design also allowed us to examine whether the condition pair with the strongest interactome proximity necessarily shared the greatest number of predicted drug-reversal candidates.

## 2. Methods

### 2.1 Data sources and differential expression analysis

Publicly available transcriptomic datasets were obtained from the NCBI Gene Expression Omnibus for each of the six skin conditions: GSE30999 (PS), GSE32924 (AD), GSE148027 (HS), GSE75819 (VL), GSE57178 (CSU), and GSE168735 (SLS). Differential expression was assessed using limma/GEO2R, comparing lesional with non-lesional skin for the five clinical conditions and SLS-exposed with control skin for SLS-ICD. DEGs were defined by an absolute log2 fold change ≥0.585 (≥1.5-fold) together with FDR-adjusted P<0.05 for HS and VL or nominal P<0.05 for PS, AD, CSU, and SLS-ICD. The nonuniform significance criteria reflect the fact that no AD or SLS-ICD genes met both the fold-change threshold and FDR<0.05 in the available datasets; consequently, all analyses that depend on DEG-set membership should be regarded as exploratory. Accession numbers, platforms, sample counts and group composition, the contrast analysed, the significance criterion applied and the resulting DEG counts are given for each dataset in Supplementary Table S2. Gene symbols in the resulting lists reflect the platform annotation available at the time of analysis. Whether the principal PS–HS and SLS-ICD findings are robust to threshold selection and to the resulting differences in DEG-set size was tested in the harmonized, equal-sized sensitivity analysis described in section 2.8.

### 2.2 Gene-level overlap analysis

Pairwise and higher-order overlap among the six DEG sets was computed as exclusive intersections (the UpSet/ComplexUpset convention): a gene is counted toward a given combination of diseases only if it is a DEG in exactly that combination and no others. This is distinct from a simple inclusive/pairwise intersection, in which a gene counts toward a pair if it is a DEG in at least both diseases regardless of its status in the remaining four; the two conventions coincide only at the most restrictive cut (all diseases sharing a gene), because there is no larger superset left to exclude against. All overlap counts reported here use the exclusive convention.

### 2.3 Gene Ontology pathway enrichment

Gene Ontology (GO) biological-process enrichment was performed for each condition using clusterProfiler.^13^ Each condition’s DEG set was mapped to Entrez identifiers and tested with compareCluster(fun = “enrichGO”, ont = “BP”) against the org.Hs.eg.db annotation, using the package defaults: the full annotated gene set as background, Benjamini–Hochberg adjustment at P < 0.05, gene sets of 10 to 500 genes, and no redundancy reduction. Up-and downregulated genes were analysed together rather than separately, so an enriched term reflects the involvement of a process and not the direction in which it changes. Because independently selected top terms can be redundant and can conceal opposite expression directions, pathway comparisons are treated as descriptive unless confirmed using direction-aware, ranked-gene enrichment.

### 2.4 Interactome-based network separation

The relative topological proximity of each condition pair was quantified using the network-separation statistic S_AB, with each DEG set mapped to the human protein–protein interaction network.^5^ This interactome-based approach was previously described by Menche et al., who constructed a map of human protein interactions comprising 13,460 proteins and 141,296 documented interactions drawn from regulatory, binary, literature-curated, metabolic, protein-complex, kinase–substrate and signaling data. In this approach S_AB<0 indicates that two modules are closer to one another than expected from their within-module distances, whereas S_AB>0 indicates relative separation. The interactome was used as published, without filtering of edges; self-interactions were removed and genes absent from the network were dropped, leaving mapped modules of 4,417 (PS), 1,075 (AD), 4,258 (HS), 2,379 (VL), 777 (CSU) and 22 (SLS-ICD) proteins. Distances are unweighted shortest paths, with d_AA taken as the mean distance from each gene in a module to its nearest other member and d_AB as the corresponding mean between modules. Empirical P values and confidence intervals from degree-and module-size-matched permutations were not computed, so S_AB is reported here as a relative descriptor rather than as an inferential test against a randomized null.

### 2.5 DisGeNET-based diseasome mapping

Each condition’s DEG set was analyzed for enrichment of curated DisGeNET disease-gene sets using disgenet2r, the HGNC vocabulary, and the curated source.^14^ Results were combined across conditions after disease-identifier harmonization. Associations meeting Benjamini–Hochberg FDR<0.25 were retained as exploratory candidates; this threshold was chosen for hypothesis generation and does not establish comorbidity.

As an additive exploration, DisGeNET was queried directly through its Model Context Protocol interface (via the DISGENET MCP Connector; *disgenet.com/Tools?section=mcp),* using an academic-tier account restricted to curated sources (CLINGEN, CLINPGX, CLINVAR, GENCC, MGD_HUMAN, ORPHANET, PSYGENET, RGD_HUMAN and UNIPROT; CTD_human was not accessible under this tier). Disease-disease associations were retrieved for each index disease by UMLS concept identifier: psoriasis C0033860, atopic dermatitis C0011615, hidradenitis suppurativa C0162836, vitiligo C0042900 and irritant contact dermatitis C0162823. This endpoint links two diseases when they share curated disease genes and applies no statistical test; a minimum of five shared genes was required for all diseases, with results for hidradenitis suppurativa additionally reported at a minimum of two, since its curated gene set of nine genes cannot support the higher threshold. No identifier corresponding specifically to chronic spontaneous or chronic idiopathic urticaria resolved in the current DisGeNET release, so CSU could not be included in this query. Gene-disease associations restricted to curated sources were retrieved separately for each of the 14 genes shared across the PS, AD, HS, VL and CSU DEG sets.

### 2.6 Real-world patient co-occurrence in TriNetX

Candidate disease associations from the DisGeNET enrichment described above were examined for patient co-occurrence in the TriNetX federated health-research network. Associations were selected at Benjamini–Hochberg FDR < 0.05 for PS, AD, HS, CSU and SLS-ICD. No association reached FDR < 0.05 for VL, whose strongest association was FDR = 0.136; for that condition the exploratory FDR < 0.25 set was used instead. Two composite DisGeNET concepts in the SLS-ICD set were split into their five component terms so that each could be mapped to an ICD-10-CM code, and two vitiligo concepts were not carried forward, yielding 220 associations for examination. Each is listed individually, with its DisGeNET FDR and its TriNetX outcome, in Supplementary Table S3.

Queries were performed manually against a single contributing healthcare organization (HCO) University of Cincinnati Medical Center, using ICD-10-CM diagnosis codes with no restriction on observation window or diagnosis type. A cohort was defined for each index condition (PS n = 10,140; AD n = 7,610; HS n = 3,490; SLS-ICD n = 1,910; VL n = 1,180; CSU n = 510). For each candidate association, the number of patients within that cohort who also carried the candidate diagnosis was recorded, together with the cohort denominator.

Patient counts returned by TriNetX are rounded, and the smallest non-zero value observed across these queries was ten. Associations returning a non-zero count are reported here as observed co-occurrences, and those returning none as not detected; an absent count cannot be distinguished from a count falling below the reporting threshold. No comparator cohorts were constructed, and no effect estimates, confidence intervals, covariate adjustment, or correction for multiple comparisons were applied.

### 2.7 Drug-repurposing analysis

Candidate compounds were identified with the signatureSearch R/Bioconductor package^15^ and LINCS L1000 Level 5 moderated z-score perturbation signatures. ^16^ For each index condition, up- and downregulated Entrez-gene signatures were defined from that condition’s DEG set and queried against the LINCS reference, which comprises 45,956 signatures representing 8,140 compounds profiled across 30 cell lines. Connectivity was scored using the weighted connectivity score implemented in signatureSearch and normalized (NCS). A negative score indicates that a compound shifts the disease-associated genes in the direction opposite to the disease itself; such profiles are designated candidate reversal signatures, whereas compounds whose profiles resemble the disease are not carried forward. Because most compounds are profiled in more than one cell line, signatures were collapsed to one entry per compound, retaining the cell line in which that compound scored most strongly; the ranks reported are therefore positions among the 7,697 compounds with a reversal-direction signature. Cross-disease comparisons used the top 50 compounds per condition, and the effect of that cut-off was examined at 25, 50, 100 and 200. The LINCS panel contains no keratinocyte-derived line; the compounds in the psoriasis top-50 list were profiled in A375, A549, ASC, HCC515, MCF7, PC3 and VCAP. A separate and nondirectional comparison was performed for the DisGeNET-predicted diseases observed to co-occur in TriNetX, which have no expression signature of their own. For each, its curated DisGeNET gene set was tested against the genes with |z|>2 in each LINCS signature using Fisher’s exact test, and the resulting top 100 compounds were compared with the top 100 reversal candidates of the corresponding skin condition. This test asks whether a compound perturbs a disease’s genes at all rather than whether it reverses them, so an overlap identified in this way indicates shared pharmacologic targeting rather than shared reversal. Post hoc, top-ranked compounds were checked against PubMed and ClinicalTrials.gov (September 2026) for existing clinical use in the corresponding skin condition or co-occurring disease.

### 2.8 Sensitivity analysis of DEG thresholds and gene-set size

To evaluate whether the use of different statistical thresholds, and the resulting variation in DEG-set size, influenced the findings, we performed complementary size-normalized and harmonized sensitivity analyses. The original analysis applied a uniform effect-size threshold of |log2FC| > 0.585 (an absolute fold change of 1.5) to all datasets, together with an FDR-adjusted p < 0.05 for HS and VL and a nominal p < 0.05 for PS, AD, CSU and SLS-ICD; a uniform FDR threshold was not feasible because no genes in AD or SLS-ICD satisfied both FDR < 0.05 and the fold-change criterion. First, pairwise similarity among the original DEG sets was normalized using the Jaccard index, defined as the number of shared genes divided by the union of the two sets. Second, an identical harmonized procedure was applied to every condition: analyses were restricted to the 10,340 genes measurable across all platforms, genes were required to meet a nominal p < 0.05, the remaining genes were ranked by descending |log2FC|, and equal-sized top-N gene sets were selected. Effect size was used for ranking rather than p-value because p-value magnitude depends on sample size, which ranged from n = 170 (PS) to n = 18 (CSU) across these datasets. Analyses including all six conditions were performed at N = 250, 400 and 582, the last being the largest set supported by SLS-ICD, whereas analyses restricted to the five clinical diseases were performed at N = 1,000, 2,000 and 2,720. SLS-ICD retained the lowest overlap with the other conditions under harmonization, at 0.47 times the mean overlap of pairs not involving it, consistently across all set sizes. Full results, including recomputed interactome separation at harmonized set sizes and comparison of the original overlaps against hypergeometric expectation, are provided in Supplementary Data S1.

## 3. Results

We organized the Results as a sequential test of concordance across disease layers. First, DEG intersections and GO enrichment defined shared and condition-specific expression programs. Second, S_AB positioned the resulting modules within the protein interactome. Third, DisGeNET expanded these modules into candidate disease associations. Fourth, selected associations were examined descriptively for clinical co-occurrence in TriNetX. Finally, LINCS L1000 signatures were used to ask whether disease pairs with similar molecular architecture also shared candidate pharmacologic reversal profiles. This sequence distinguishes discovery, contextualization, clinical observation, and therapeutic hypothesis generation.

### 3.1 Gene-level overlap among the six skin diseases

The largest exclusive DEG intersection was observed between PS and HS (2,060 genes), followed by AD–PS–HS (490 genes), VL–PS (464 genes), AD–PS (402 genes), and VL–PS–HS (379 genes) (Figure 1). Fourteen genes—CXCL8, STAT1, LYZ, CTSC, SRGN, NMI, CORO1A, JAML, PTGS2, DENND4A, FHL1, SELENOP (SEPP1), KRT15, and SSBP2—were present in the DEG sets of PS, AD, HS, VL, and CSU but not SLS-ICD. These genes span neutrophil recruitment, interferon signaling, lysosomal and antimicrobial functions, immune-cell organization, prostaglandin synthesis, epithelial biology, and transcriptional regulation. Because exclusive counts depend on the sizes and thresholds of the contributing DEG sets, they are reported here as descriptive. Size-normalized overlap and comparison against hypergeometric expectation, which address whether PS–HS sharing exceeds that of other pairs once set size is taken into account, are presented in section 3.4 and Supplementary Data S1.

**Figure 1.**
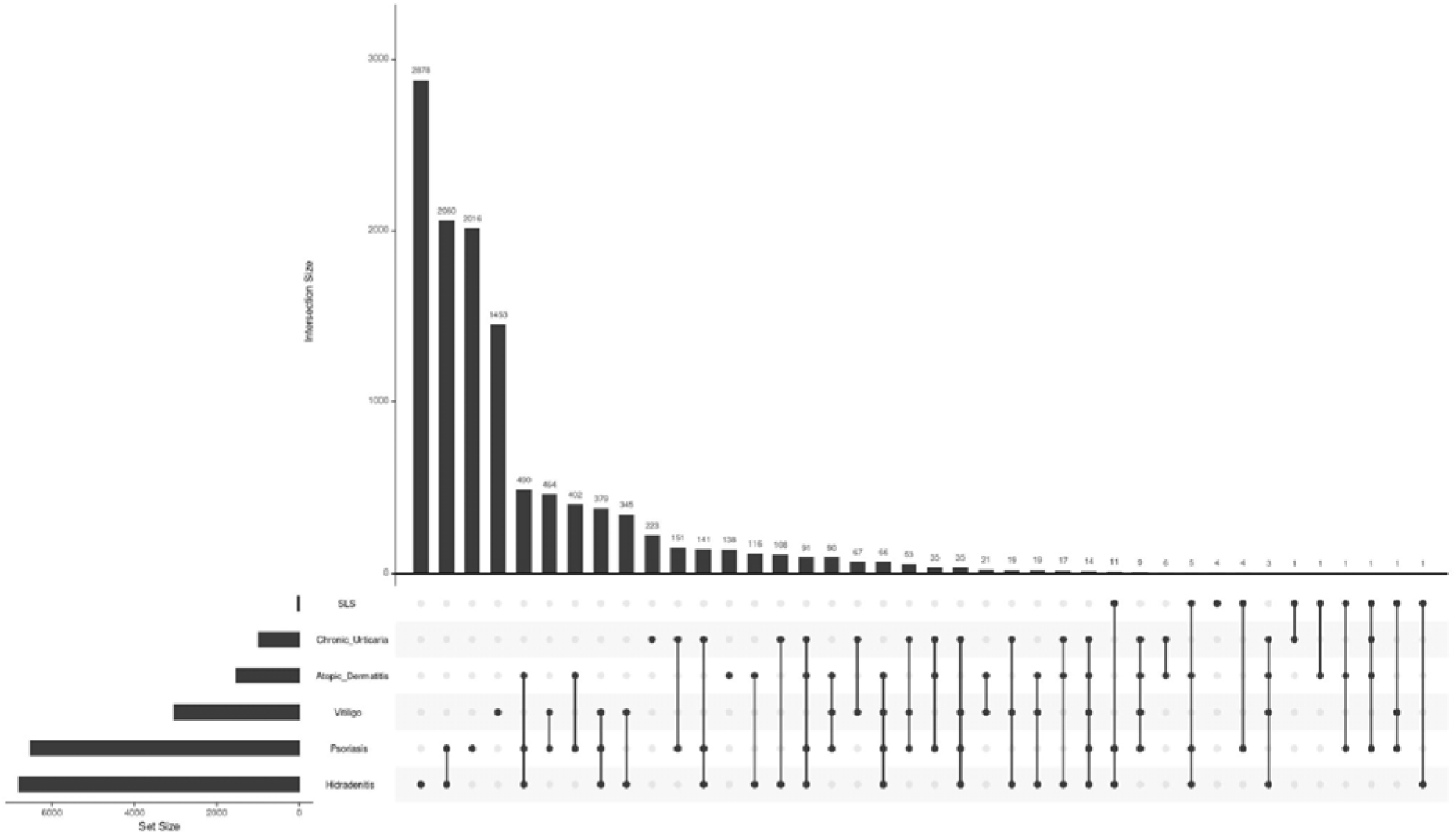
Exclusive differentially-expressed-gene (DEG) overlap among the six skin diseases (UpSet plot).

### 3.2 Pathway-level overlap

GO biological-process enrichment revealed recurrent neutrophil activation and neutrophil-mediated immunity among PS, HS, and CSU, and T-cell activation among AD and HS (Figure 2). AD additionally showed lipid and fatty-acid metabolic programs. VL was characterized among its leading terms by ribosome biogenesis and RNA metabolism, whereas SLS-ICD was dominated by keratinization, cornification, epidermal development, and skin development. The SLS-ICD profile is consistent with a chemically induced barrier-injury model. These comparisons describe the leading enriched terms within each condition. They do not establish that a pathway is absent when it is not top ranked, and because up- and downregulated genes were tested together, a term shared by two conditions does not imply that it changes in the same direction in both.

**Figure 2.**
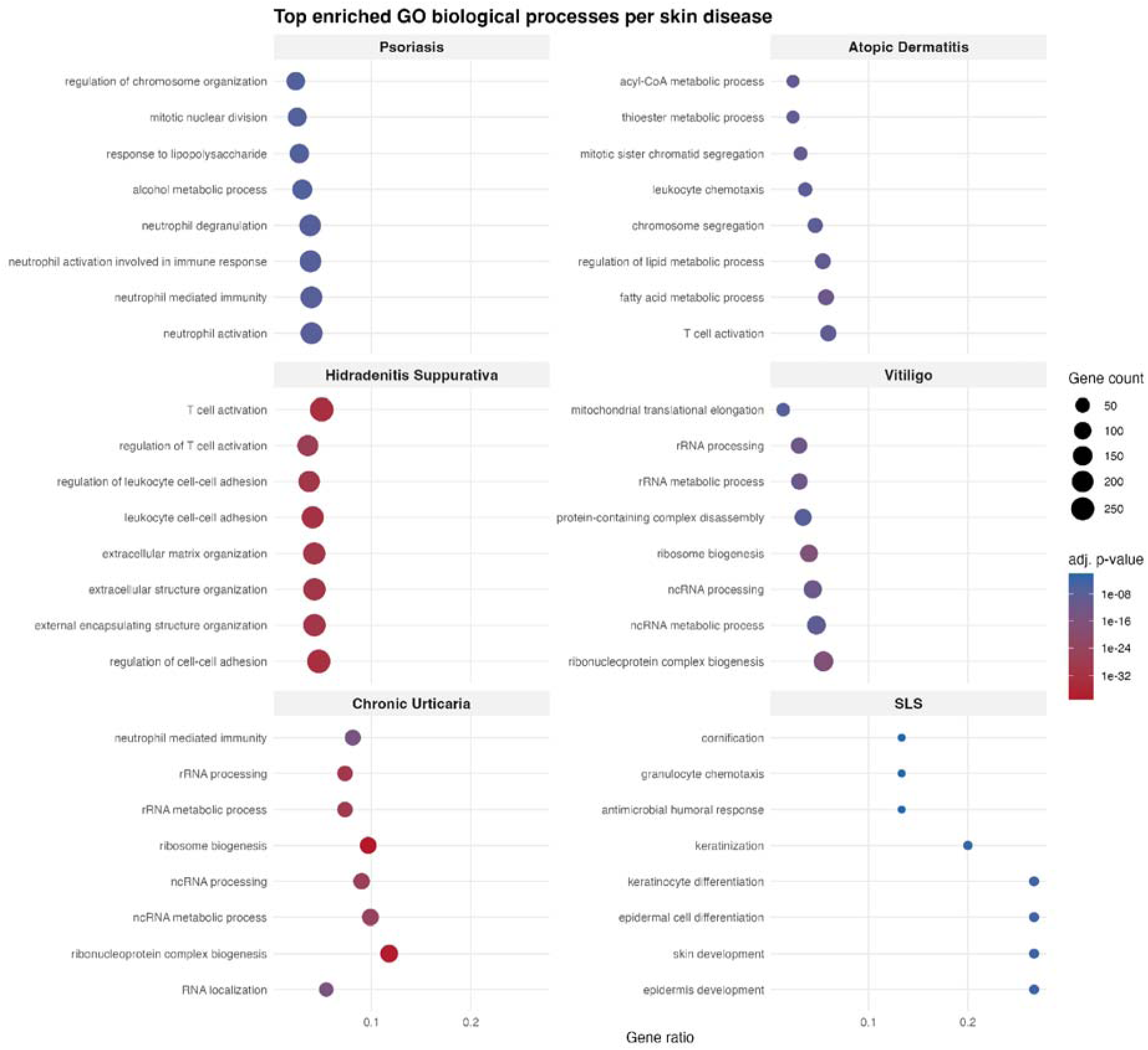
Top enriched Gene Ontology biological process terms per disease.

### 3.3 Network-based molecular relatedness

Observed S_AB values were lowest for PS–HS (−0.594), followed by PS–VL (−0.272), AD–PS (−0.267), HS–VL (−0.185), AD–HS (−0.152), and AD–CSU (−0.132) (Figure 3). Eight of the ten pairings among PS, AD, HS, VL, and CSU had negative point estimates. The two exceptions both involved CSU and lay close to zero (PS–CSU 0.020, HS–CSU 0.050), placing CSU at the margin of the immune-mediated group rather than within it. In contrast, SLS-ICD had positive values with HS (0.608), PS (0.597), VL (0.507), AD (0.418), and CSU (0.332), indicating relative topological separation from the five immune-mediated conditions. This pattern is concordant with the distinct SLS-ICD pathway profile. Because empirical uncertainty was not estimated in the current analysis, these values are interpreted as relative topological descriptors rather than statistically significant departures from a randomized null.

**Figure 3.**
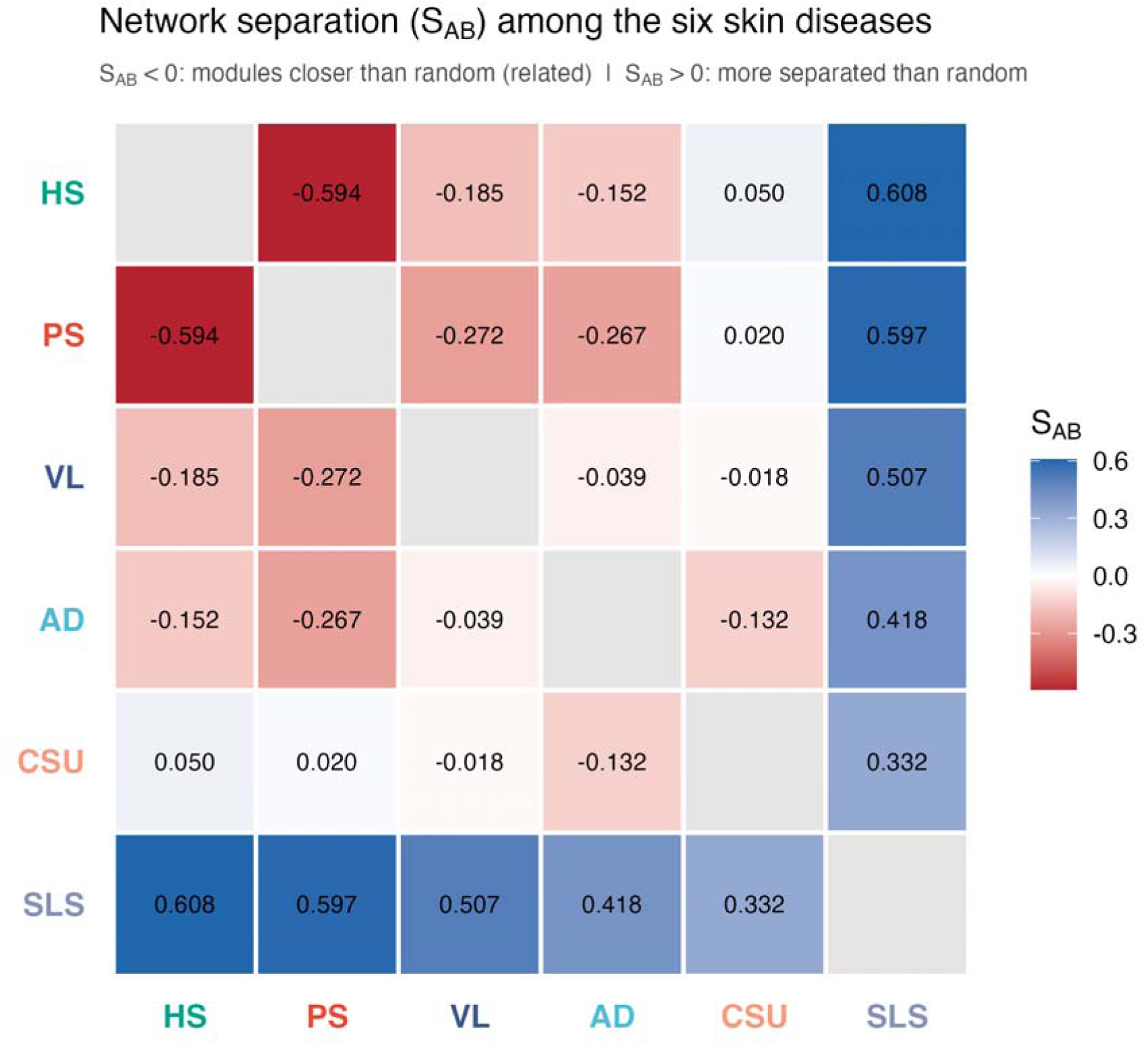
Interactome-based network separation (S_AB) among the six skin diseases

### 3.4 Sensitivity analysis of DEG thresholds and module size

To determine whether differences in statistical thresholds and DEG-set sizes influenced the results, we performed size-normalized and harmonized sensitivity analyses. Using the complete DEG lists, PS–HS remained the strongest pair after normalization by the Jaccard index (0.329), followed by PS–AD (0.176), indicating that the large PS–HS overlap was not explained solely by gene-set size. We next applied an identical procedure across datasets by restricting the analysis to genes measurable across platforms, requiring nominal P < 0.05, ranking genes by |log2 fold change|, and comparing equal-sized top-N sets. PS–AD and PS–HS consistently emerged as the two strongest pairs across all evaluated set sizes, although their relative ranking depended on analysis depth: PS–AD predominated among the most highly ranked genes, whereas PS–HS led when the complete DEG lists were considered. All ten pairings among PS, AD, HS, VL, and CSU returned negative S_AB values at both harmonized set sizes, including the two CSU pairings that were marginally positive in the primary analysis. Their relative interactome proximity was therefore robust to threshold selection and module-size equalization, and CSU’s marginal separation in the primary analysis is attributable to the smaller size of its module rather than to its position in the interactome. In contrast, the positive values originally observed for SLS-ICD were not maintained after equalization, indicating that its apparent topological separation was strongly influenced by its small DEG set. Accordingly, SLS was not interpreted as an unusually disconnected interactome module. Its distinction from the five clinical diseases was instead supported at the pathway level, where SLS showed enrichment for keratinocyte differentiation, cornification, and barrier-associated programs rather than the predominantly immune-activation programs observed in the clinical conditions.

### 3.5 DisGeNET-based diseasome mapping

At exploratory FDR<0.25, DisGeNET enrichment returned 33 candidate associations for PS, 145 for AD, 136 for HS, 12 for VL, 749 for CSU, and 259 for SLS-ICD. The leading disease classes differed across conditions and included immune, cardiovascular, neurologic, digestive, congenital/hereditary, metabolic, and neoplastic categories (Figure 4). These raw counts are not directly comparable because they reflect DEG-set size, annotation coverage, and the permissive discovery threshold. Figure 4 should therefore be interpreted as a map of candidate links rather than a quantitative comorbidity burden. The associations carried forward to the subsequent layers were those that recurred across evidence types, and the criteria used to select them are given in sections 2.6 and 2.7.

**Figure 4.**
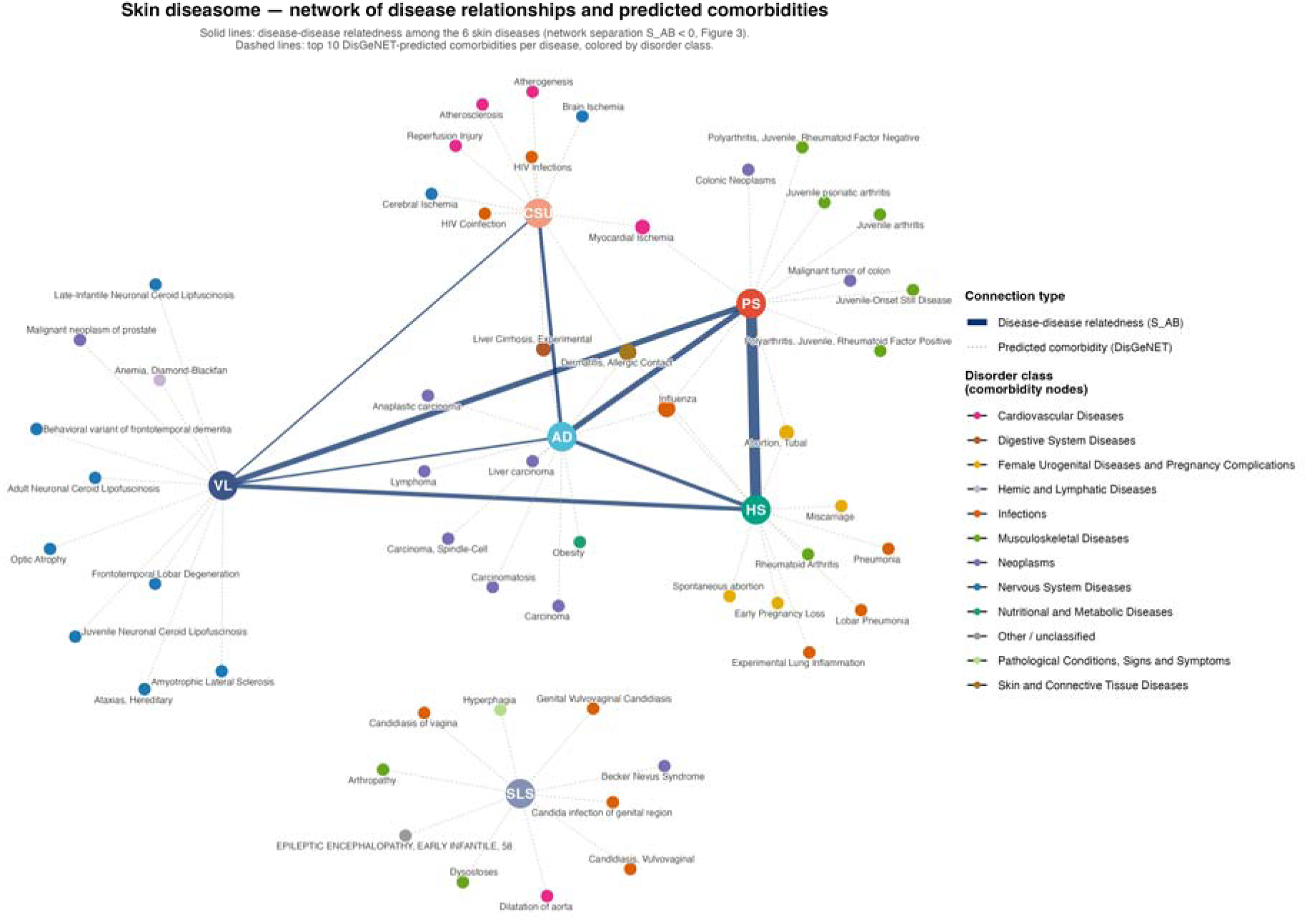
Skin diseasome network: S_AB disease-disease relatedness (solid lines) combined with top DisGeNET-predicted comorbidities (dashed lines, colored by disorder class).

### 3.6 Additive exploration of disease-disease bridging and the 14 shared genes

The above analysis captured diseases in general whose known disease-associated genes overlapped with DEGs in each skin condition. Because this approach is restricted to genes upregulated or downregulated in lesional skin, it may miss diseases that share inherited susceptibility genes with a skin disease when those genes are not differentially expressed in the skin. To examine this complementary form of relationship, we directly queried DisGeNET’s curated records. We asked two questions: (1) which diseases are documented in DisGeNET as sharing disease-associated genes with each of the six skin conditions, and (2) which diseases are individually associated with each of the 14 genes shared by PS, AD, HS, VL, and CSU.

Disease-disease associations were returned for four of the six index diseases. PS was associated with 63 other conditions, the strongest being Graves disease (16 shared genes, Jaccard 0.28), Behçet syndrome (13, 0.22), Sjögren’s syndrome (11, 0.26), systemic lupus erythematosus (14, 0.17) and psoriatic arthritis (8, 0.22), alongside rheumatoid arthritis, ankylosing spondylitis, autoimmune thyroiditis, type 1 diabetes, Crohn’s disease, multiple sclerosis, atherosclerosis, myocardial infarction and heart failure. AD was associated with 40 conditions, headed by asthma (22 shared genes, its single strongest association), atopic rhinitis, respiratory allergy, chronic obstructive airway disease, allergic contact dermatitis and Behçet syndrome. VL was associated with 24 conditions, headed by autoimmune thyroiditis (7, 0.27), Behçet syndrome (8, 0.16), Graves disease (8, 0.15), systemic lupus erythematosus (8, 0.10) and melanoma (6, 0.12), together with Sjögren’s syndrome, multiple sclerosis, rheumatoid arthritis, systemic scleroderma and hepatitis B and C. HS returned no associations at this threshold, which its curated gene set of nine genes cannot support; lowering the requirement to two shared genes returned three associations, all dermatological and all with the highest similarity values observed anywhere in this analysis: acne, acne varioliformis and pyoderma gangrenosum (4 shared genes each, Jaccard 0.80). SLS returned no associations at either threshold. CSU could not be included: no chronic spontaneous or idiopathic urticaria concept resolves in the current DisGeNET release, and the general urticaria concepts available are not specific to it.

At the individual-gene level, the 14 shared genes showed markedly heterogeneous disease association profiles. *PTGS2* (COX-2) was the most pleiotropic, with associations spanning cancers, cardiovascular disease, metabolic disease, and neuropsychiatric conditions. *STAT1* showed a concentrated immune/infection association profile, including monogenic immunodeficiencies and autoimmune disease. *CXCL8* showed a notable psychiatric/mood-disorder association cluster (delirium, bipolar disorder, depression), which is independently corroborated by clinical studies of bipolar disorder patients showing elevated serum interleukin-8 and altered peripheral-blood CXCL8 gene expression.^17,18^ *SSBP2* also returned a bipolar disorder association in DisGeNET, but we could not identify independent literature specifically linking *SSBP2* to bipolar disorder beyond this database entry, which most likely reflects a nearest-gene annotation from a psychiatric genome-wide association study locus rather than a functionally established relationship; we report this association but do not treat it as independently corroborated. *CTSC* and *FHL1* returned only high-confidence Mendelian disease associations specific to those genes (Papillon-Lefèvre disease and neuromuscular/cardiac myopathies, respectively), and *CORO1A* and *LYZ* returned only their own causal monogenic disease. *SRGN, NMI, JAML, DENND4A,* and *SEPP1* returned no disease associations under the curated data sources accessible for this analysis.

### 3.7 Descriptive patient co-occurrence in TriNetX

Of 220 DisGeNET-predicted disease associations checked against real-world patient data via TriNetX, 55 (spanning 42 distinct diseases) were corroborated by a non-zero patient co-occurrence ratio; the remaining 165 were not detected in this cohort. Corroborated associations included well-established comorbidities such as rheumatoid arthritis, atherosclerosis, asthma, and HIV infection, alongside several conditions, including juvenile (psoriatic) arthritis and allergic contact dermatitis, that were independently corroborated as related to more than one of the six index skin diseases. The complete set of queried associations and their outcomes is provided in Supplementary Table S3.

### 3.8 Drug-repurposing analysis

Matching each disease’s DEG signature against the LINCS L1000 reference identified mTOR/PI3K/Aurora-kinase pathway inhibitors (including AZD-8055, KU-0063794, and GSK-1070916) as the dominant class of top-ranked reversal candidates for PS, consistent with the established role of PI3K-AKT-mTOR signaling in keratinocyte hyperproliferation in psoriatic disease. Protein kinase C (PKC)-family inhibitors were also represented among PS’s top candidates, including midostaurin (historically developed as the PKC inhibitor PKC412; rank 43 of the 7,697 compounds with a reversal-direction signature) and the PKC-β-selective inhibitor CGP-53353 (rank 580).

Among the top 50 reversal candidates, taken as one entry per compound, PS and CSU shared 19 compounds, whereas PS and HS shared two, despite PS–HS having the lowest observed S_AB value (Figure 5A). This contrast should be interpreted cautiously, because no HS reversal signature reached FDR < 0.05 (lowest FDR = 0.31), whereas significant signatures were obtained for every other condition. This ordering did not depend on the depth of the comparison: PS–CSU was the most overlapping of the fifteen pairs at every cut-off examined (7 shared compounds in the top 25, 19 in the top 50, 48 in the top 100 and 110 in the top 200), while PS–HS shared 0, 2, 4 and 16 respectively. Across the 15 condition pairs, S_AB and shared-compound count were weakly inversely correlated (Spearman ρ=−0.22, P=0.434), an association that was not statistically significant. PS and CSU illustrate the discordance most directly: their disease modules were marginally separated in the interactome (S_AB = 0.020), yet they shared more top-ranked compounds than any other pair. Moreover, the 15 pairwise observations are not independent because each condition contributes to multiple pairs. The result is therefore reported as a descriptive discordance between interactome topology and drug-signature overlap; a dyadic permutation test and a chance-adjusted measure of compound overlap, which would allow formal inference, were not performed. All skin condition–comorbidity pairs for which this comparison was performed, together with the shared candidate compounds for each, are provided in Supplementary Table S1.

**Figure 5.**
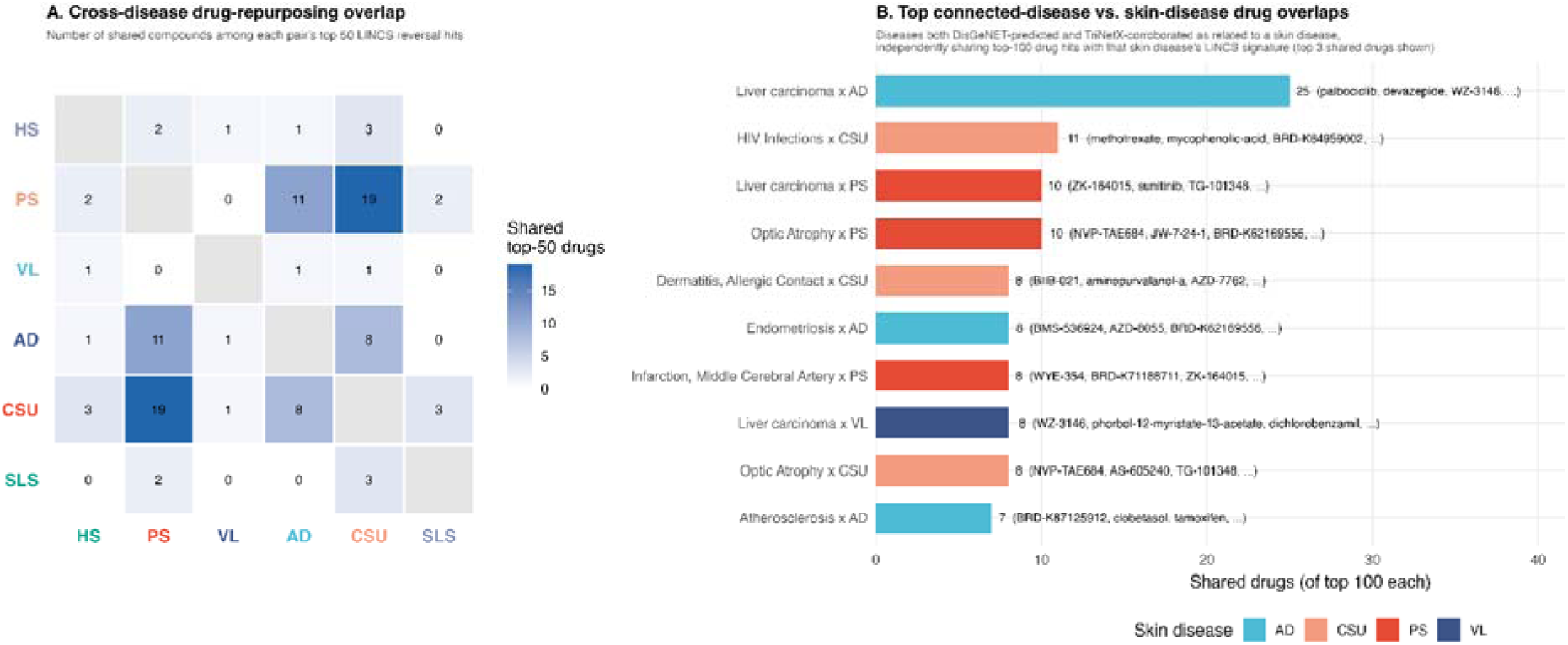
Cross-disease drug-repurposing overlap and top connected-disease vs. skin-disease drug overlaps.

Comparison of each skin condition’s top 100 reversal candidates with the nondirectionally derived compound lists of selected DisGeNET-linked diseases observed to co-occur in TriNetX identified several overlaps, including 25 compounds between AD and liver carcinoma and 11 between CSU and HIV infection (Figure 5B). Interestingly, methotrexate and mycophenolate mofetil appeared in both the CSU-derived and HIV-derived candidate lists and have prior use in chronic or refractory urticaria.^33,34^ This recovery of clinically familiar immunosuppressants supports the face validity of the prioritization strategy, but it does not constitute independent validation: both compounds may emerge through broad antiproliferative or immunosuppressive signatures. Predictive performance was not quantified: the analysis was not benchmarked against known-indication sets or against unrelated immune-disease controls. Several top-ranked candidates are already in clinical use. For their own condition, topical corticosteroids ranked highly for AD (clocortolone pivalate 3rd and mometasone 10th of 7,869)^19^, sirolimus^20^ for PS (36th of 7,697) and mycophenolate mofetil for CSU^21^ (10th of 7,657), although first-line H1-antihistamines for CSU ranked beyond 1,000th. Compounds linking a skin condition to a co-occurring diseases were also used or tested in both including methotrexate^37^ , statins (lovastatin/ pitavastatin in CSU and HIV-associated cardiovascular inflammation)^22,23^ and sirolimus (PS and infection)^20,24^. Among candidates without such precedent, midostaurin ranked 7th for CSU (FDR < 0.001). It inhibits KIT, which sustains mast cells, the principal effector cells of CSU, and is approved for systemic mastocytosis.^25^ KIT-targeted mast-cell depletion has also shown clinical activity in CSU.^26^

## 4. Discussion

Collectively, three major findings define the central story of this study. First, the five immune-mediated diseases formed a connected core across the DEG, pathway and interactome layers, with PS and HS showing the greatest observed topological proximity; CSU sat at the margin of that core, its interactome separation from PS and HS being marginally positive in the primary analysis and negative once module sizes were harmonized. Second, SLS-ICD remained distinct at the transcriptomic and pathway levels, consistent with its role as an experimentally induced barrier-injury comparator, although its apparent interactome separation did not survive harmonization of gene-set size. Third, pharmacologic reversal overlap did not mirror interactome proximity: PS and CSU, which were marginally separated in the interactome, shared more top-ranked candidates than any other pair. Together, these findings indicate that disease relatedness is multidimensional and that molecular topology cannot be used as a proxy for shared therapeutic response.

The PS–HS relationship provides the clearest example of cross-layer convergence. Its large exclusive DEG intersection, shared immune pathways, lowest observed S_AB value, and established epidemiologic association all point toward overlapping inflammatory architecture^19,20^ . In contrast, the PS–CSU drug-signature overlap illustrates cross-layer divergence. Interactome topology reflects the organization of disease-associated proteins, whereas LINCS reversal scores reflect context-dependent perturbational responses in specific cell lines, doses, and exposure times. These layers can therefore agree when diseases share both network neighborhoods and reversible expression programs, but they need not do so.

These molecular relationships are broadly consistent with the clinical literature. Although psoriasis (PS), atopic dermatitis (AD), hidradenitis suppurativa (HS), vitiligo (VL), and chronic spontaneous urticaria (CSU) are clinically distinct disorders, epidemiologic studies increasingly demonstrate that they can co-occur within the same individuals, suggesting overlapping immune and genetic susceptibility. The strongest evidence among the conditions examined here is for the association between PS and HS. A large population-based study found that HS occurred more frequently among patients with PS than matched controls (0.3% vs. 0.2%; adjusted OR, 1.8), and a subsequent systematic review and meta-analysis further supported an increased risk of PS among individuals with HS.^27,28^ AD and HS also show a bidirectional relationship: patients with HS had approximately twice the risk of developing AD compared with matched controls (HR, 2.06), while pre-existing AD was more prevalent among patients with HS.^29^ AD has additionally been associated with VL; a systematic review and meta-analysis of more than 1.5 million patients with AD found increased odds of VL (OR, 3.21) and an increased risk in cohort studies (RR, 1.64).^30^ PS and VL have likewise been reported to co-occur, although the strength and consistency of this association appear weaker than those observed for PS–HS or AD–VL.^31,32^ CSU is also characterized by substantial autoimmune comorbidity and has been repeatedly associated with PS and VL. A systematic review found that PS occurred more frequently among patients with CSU and that VL was present in approximately 3% of patients, with autoimmune thyroid disease and other autoimmune disorders occurring particularly frequently.^33^ Thus, the clinical literature supports a partially interconnected network of inflammatory and immune-mediated skin diseases rather than six completely independent disease entities.

The inclusion of sodium lauryl sulfate (SLS) in this framework requires a different interpretation. SLS is not a clinical disease entity but a widely used experimental model of irritant contact dermatitis (ICD), producing controlled epidermal injury, inflammation, and barrier dysfunction.^34^ Unlike PS, AD, HS, VL, and CSU, there is therefore no meaningful epidemiologic literature establishing SLS “comorbidities.” Nevertheless, ICD and AD can coexist clinically, and impaired barrier function may increase susceptibility to irritant and contact reactions^35^. Accordingly, SLS-induced dermatitis in the present study should be viewed as a mechanistically informative barrier-injury comparator rather than as a disease with established clinical comorbidity relationships. This distinction is important because it allows molecular overlap with SLS to be interpreted as evidence of shared pathways associated with epithelial injury, barrier disruption, or inflammation, rather than as evidence of a direct clinical disease association.

SLS-ICD was a coherent negative comparator: it did not share the 14-gene signature found across the five immune-mediated diseases, its leading pathways emphasized keratinization and epidermal differentiation, and its S_AB point estimates were positive in all five comparisons. This convergence is biologically plausible, but it should not be treated as definitive internal validation. SLS-ICD differs from the clinical datasets in exposure, sampling design, and disease context, and some separation may therefore reflect study architecture or DEG-set construction. The harmonized analysis reported in Supplementary Data S1 bears directly on this point. When every condition was required to contribute an identically sized gene set, the pathway distinction and the comparatively low gene overlap of SLS-ICD persisted, whereas its positive S_AB values did not, indicating that the apparent topological separation was governed by the small size of the SLS-ICD module rather than by its position in the interactome. We therefore rest the SLS-ICD distinction on the pathway and overlap evidence rather than on interactome topology.

The direct DisGeNET queries recovered, from curated disease genetics alone, much of the comorbidity profile these diseases are known for clinically. PS was linked to psoriatic arthritis, ankylosing spondylitis, Crohn’s disease, atherosclerosis and cardiac disease. AD’s strongest single association was asthma, the defining transition of the atopic march. VL’s were dominated by autoimmune thyroiditis, Graves disease, Behçet syndrome, systemic lupus erythematosus and Sjögren’s syndrome. HS is curated with only nine genes, yet those genes are shared almost entirely with acne and pyoderma gangrenosum, the conditions with which it co-occurs in the follicular occlusion and PASH syndromes, and the most specific association returned anywhere in this analysis. SLS returned no disease–disease associations at either threshold.

These associations derive predominantly from curated model-organism orthology, supplemented by variant-level curation; the human GWAS-derived sources sit outside the access tier used here. They therefore corroborate clinical observation rather than supply independent human genetic evidence. Two boundaries are worth naming: chronic spontaneous urticaria has no matching concept in the current DisGeNET release and could not be queried, and SSBP2’s bipolar-disorder association has no support beyond the database entry, most likely a nearest-gene annotation from a psychiatric GWAS locus. CSU’s absence here is a terminology gap rather than a biological one; the CSU comorbidities identified via the TriNetX-corroborated drug-repurposing route show that CSU does have detectable real-world disease associations. The remaining gene-level results are consistent and interpretable. PTGS2’s pleiotropy and STAT1’s concentration in immune and infection phenotypes match their established roles, and CXCL8’s mood-disorder cluster is independently supported by clinical studies showing elevated serum interleukin-8 and altered peripheral-blood CXCL8 expression in bipolar disorder.^17,18^

In the descriptive TriNetX layer, 55 of 220 queried associations, representing 42 distinct candidate diseases, had non-zero co-occurrence with an index condition; 165 were not detected. These results indicate presence or absence within the queried cohorts, not enrichment relative to controls. Non-detection may reflect cohort composition, coding, query definitions, or privacy-related count suppression; associations returning no count are therefore reported as not detected rather than refuted, and the observed subset represents a lower bound on real-world relevance rather than an exhaustive account of it. Accordingly, the observed subset should be regarded as a prioritization set for controlled epidemiologic analysis rather than a validated comorbidity network.

The drug-repurposing findings illustrate both the promise and the limits of this kind of computational prediction. The dominant reversal signal for psoriasis, mTOR/PI3K/Aurora-kinase pathway inhibitors, corroborates, rather than newly discovers, a therapeutic hypothesis already present in the literature: PI3K-AKT-mTOR signaling is upregulated in psoriatic skin and implicated in keratinocyte hyperproliferation, and mTOR inhibition has separately been proposed and tested as an anti-psoriatic strategy.^36–38^ Together with the recovery of established treatments, including drugs used in both a skin condition and its co-occurring disease, this supports the face validity of the ranking, although the check was post hoc and did not recover antihistamines for CSU. The PKC-family signal for psoriasis (midostaurin, CGP-53353) is more tentative: midostaurin’s mechanism as a PKC inhibitor is well established^39^, and it has separate precedent as a repurposed dermatological therapeutic in a different autoimmune blistering skin disease, pemphigus^40^, but we are not aware of any study testing midostaurin specifically in psoriasis; this should be regarded as a computationally generated hypothesis requiring experimental follow-up, not an established finding. Midostaurin is a stronger candidate for CSU, where it ranked 7th and its KIT target has been clinically validated by mast-cell–depleting therapy [PMID:40415544].

The recovery of methotrexate and mycophenolate mofetil from both CSU- and HIV-derived signatures provides a clinically recognizable proof of concept, because both drugs have been used in chronic or refractory urticaria. ^41,42^ Nevertheless, this convergence should be interpreted as hypothesis support rather than validation. The result may reflect a shared lymphocyte-proliferation or inflammatory program, but it may also arise from nonspecific antiproliferative effects that occur across immune-disease signatures.

One plausible explanation is convergence on activated lymphocyte programs. Chronic HIV infection can involve polyclonal B-cell activation, hypergammaglobulinemia, and autoantibody production, while a subset of CSU is autoimmune and involves IgG directed against IgE or FcεRI.35 Mycophenolate inhibits de novo purine synthesis in proliferating lymphocytes,36 and low-dose methotrexate has been studied as an anti-inflammatory intervention in treated HIV infection, although ACTG A5314 did not improve endothelial function or inflammatory biomarkers and produced more adverse events than placebo.37 These observations make the CSU–HIV link biologically plausible but not specific. Enrichment of the shared drug signal for B-cell/autoantibody pathways relative to matched unrelated immune-disease pairs would be needed to distinguish a pathway-specific relationship from generalized immunosuppression.

This study integrates several complementary evidence layers and generates testable hypotheses, but its limitations are substantial. First, the source transcriptomic datasets differ in sample size, tissue and lesion context, platform, patient composition, and statistical power; cross-study batch effects and cell-composition differences may therefore contribute to apparent disease relationships. Second, DEG definitions were not statistically uniform: HS and VL used FDR-adjusted thresholds, whereas PS, AD, CSU, and SLS-ICD used nominal P values. Because every downstream layer depends on DEG-set membership and size, we performed harmonized-threshold and equal-sized sensitivity analyses (Supplementary Data S1); these confirmed the interactome proximity of the five clinical conditions but showed that the identity of the single closest pair, and the apparent separation of SLS-ICD, are sensitive to those choices. Third, exclusive intersections and the 15 pairwise comparisons are non-independent; size-normalized (Jaccard) and hypergeometric comparisons are reported in Supplementary Data S1, but permutation-based inference was not performed. Fourth, the interactome is incomplete and context-free, and uncertainty around S_AB was not estimated.

Fifth, the disease–disease associations retrieved from DisGeNET derive predominantly from mouse and rat model-organism databases (MGD_HUMAN, RGD_HUMAN), as the human GWAS-derived sources were not accessible under the academic-tier licence used here; these associations therefore reflect orthology-based inference rather than direct human genetic evidence.

Sixth, DisGeNET coverage and annotation density vary across conditions, and the FDR<0.25 layer is explicitly exploratory. Seventh, the TriNetX component was a descriptive presence/absence query performed against a single contributing healthcare organization, without matched comparator cohorts, adjusted effect estimates or confidence intervals, adjustment for age, sex, race and ethnicity, healthcare utilization or treatment exposure, or correction for multiple comparisons; counts are additionally subject to a minimum reporting threshold. It therefore prioritizes candidate associations for controlled testing and cannot validate comorbidity, which would require a prespecified matched-cohort design. Eighth, LINCS signatures may not reproduce skin-relevant cell states, exposures, or pharmacology; top-ranked compounds may reflect generalized toxicity or antiproliferative effects, and none was experimentally tested here. Finally, SLS-ICD represents a single experimental perturbation rather than a clinical cohort, limiting direct comparability. These limitations constrain causal and therapeutic interpretation but define a clear validation pathway: permutation-based inference on the network layer, controlled clinical association testing, and mechanistic evaluation in relevant skin models.

Taken together, this study maps a structured but non-uniform skin diseasome in which transcriptomic overlap, pathway convergence, interactome topology, clinical co-occurrence, and drug-reversal signatures provide complementary evidence. The five immune-mediated diseases formed a connected molecular core, SLS-ICD served as a distinct barrier-injury comparator, and predicted therapeutic overlap was partially decoupled from network proximity. The principal contribution is therefore not a claim of validated comorbidity or efficacy, but a transparent framework for prioritizing disease relationships and candidate interventions for controlled epidemiologic and experimental testing.

## Data and Code Availability

All transcriptomic datasets analysed in this study are publicly available from the NCBI Gene Expression Omnibus under accessions GSE30999 (PS), GSE32924 (AD), GSE148027 (HS), GSE75819 (VL), GSE57178 (CSU) and GSE168735 (SLS-ICD). Processed differential-expression tables, the harmonized and ranked gene sets and tested-gene universes used in the sensitivity analysis, GO enrichment outputs, the protein-protein interaction network and per-disease gene lists used for the network-separation calculation, the DisGeNET enrichment table underlying the diseasome, aggregate TriNetX co-occurrence counts with the cohort definitions used to generate them, compound-level LINCS L1000 rankings for all six conditions, and the analysis scripts that generate every figure and table reported here are available at https://github.com/Saadman/skin_diseasome. Package versions and the computing environment are recorded in SESSION_INFO.txt in that repository. Individual-level TriNetX records cannot be redistributed under the terms of the platform licence; the aggregate counts reported here are provided in full. Use of the DisGeNET API requires a licence from the provider, and the LINCS L1000 reference is distributed through Bioconductor ExperimentHub; the query outputs derived from both are included in the repository.

## Supporting information

Supplementary_Data_S1_sensitivity_analysis

Supplementary_Table_S1_drug_candidates

Supplementary_Table_S3_trinetx_associations

