## Supplementary_Data_S1_sensitivity_analysis for "A Multilayer Human Skin Diseasome Links Molecular Architecture, Clinical Co-occurrence, and Drug-Reversal Signatures"

### Supplementary Data S1. Sensitivity of the reported findings to differing statistical thresholds

Gene-set size influences overlap counts, enrichment results and interactome module size, and the six conditions analysed here were not thresholded identically. This supplement reports sensitivity analyses testing whether the findings depend on those choices: first by re-expressing the reported overlaps in a size-normalised form, then by repeating the analysis with an identical rule and equal set sizes applied to every condition. Section 4 addresses SLS-ICD separately, since its gene set is the smallest and the most affected by these considerations.

#### 1. What thresholds were used, and why they differ

A uniform fold-change criterion of $|log2FC| > 0.585$ was applied to all six datasets. The significance criterion differs: an FDR-adjusted $p < 0.05$ for HS and VL, and a nominal $p < 0.05$ for PS, AD, CSU and SLS-ICD. This was not a preference. The table below shows the number of genes each rule yields.

The criterion applied to each disease can be read directly from its gene list. A list thresholded at FDR < 0.05 cannot contain a gene whose adjusted p-value exceeds 0.05. The table reports the largest nominal and adjusted p-value present in each published list.

| Disease | Genes analysed | Largest nominal p in list | Largest adjusted p in list | Criterion |
|---|---|---|---|---|
| PS | 6,522 | 0.0495 | **0.102** | nominal p < 0.05 |
| AD | 1,527 | 0.0500 | **0.507** | nominal p < 0.05 |
| CSU | 976 | 0.0487 | **0.229** | nominal p < 0.05 |
| SLS-ICD | 33 | 0.0481 | **0.974** | nominal p < 0.05 |
| VL | 3,040 | 0.011 | **0.0499** | FDR < 0.05 |
| HS | 6,801 | 0.0259 | **0.051** | FDR < 0.05 |

PS, AD, CSU and SLS-ICD each contain genes with adjusted p-values between 0.10 and 0.97, which no FDR-based cut could admit, identifying them as nominally thresholded. Their largest nominal p-values also sit immediately below the cut (0.0481 to 0.0500), as expected when the nominal threshold is the binding constraint.

VL and HS show the opposite pattern. Their adjusted p-values reach the FDR boundary (0.0499 and 0.051 respectively, with 6,800 of 6,801 HS genes below 0.05), while their largest nominal p-values stop well short of 0.05 (0.011 and 0.0259), because for these two diseases the FDR criterion bound first. A uniform fold-change criterion of $|log2FC| > 0.585$ was applied to all six diseases.

A uniform FDR threshold across all six diseases is not available. Applying FDR < 0.05 together with the fold-change criterion to the underlying differential-expression output removes every gene in AD and in SLS-ICD, which would eliminate two diseases from the study. The per-disease criteria are therefore the only ones under which all six diseases can be analysed.

#### 2. Size-normalised overlap of the reported gene sets

The concern is that larger gene sets overlap more. This can be tested directly on the published sets using the Jaccard index, which divides shared genes by the union of the two sets and therefore removes the influence of set size. If the PS-HS result were an artefact of PS and HS being the two largest sets, it would not survive this correction.

| Disease pair | Shared genes | Jaccard index |
|---|---|---|
| **PS-HS** | **3,295** | **0.329** |
| PS-AD | 1,206 | 0.176 |
| PS-VL | 1,113 | 0.132 |
| AD-HS | 823 | 0.110 |
| HS-VL | 882 | 0.098 |
| PS-CSU | 532 | 0.076 |

PS-HS remains the strongest pair after normalisation, by a margin of nearly two-fold over the next pair. The ranking is therefore not produced by set size alone.

#### 3. Harmonised analysis: identical rules and equal set sizes

To remove the threshold and the size difference simultaneously, one identical rule was applied to every disease: (i) restrict to genes measurable on all platforms; (ii) require nominal $p < 0.05$; (iii) rank the surviving genes by |log2FC| and take the top N, with the same N for every disease.

Ranking uses effect size rather than p-value because p-value depends on sample size (PS $n = 170$, CSU $n = 18$), which would reintroduce the imbalance being controlled for. The significance filter is applied first, because ranking on effect size alone selects genes that are not distinguishable from noise.

N is limited to 582 when all six diseases are included, this being the maximum SLS-ICD can supply. Excluding SLS-ICD, the five clinical diseases support N up to 2,720, and both are reported.

##### 3a. Gene overlap under harmonisation

| Set size N | Strongest pair | PS-HS rank | PS-AD vs PS-HS |
|---|---|---|---|
| 250 (six diseases) | PS-AD | 2nd of 15 | 71 vs 26 |

| | | | |
|---|---|---|---|
| 582 (six diseases) | PS-AD | 2nd of 15 | 202 vs 99 |
| 1,000 (five diseases) | PS-AD | 2nd of 10 | 361 vs 242 |
| 2,720 (five diseases) | PS-AD | 2nd of 10 | **1,101 vs 1,086** |

PS-AD and PS-HS are consistently the two strongest pairs. Their order depends on how far into each ranked list the comparison extends: PS-AD leads at the top of the list, while the difference narrows to 1.4 per cent at the deepest level tested, and PS-HS leads when the complete gene lists are used (Section 2).

##### 3b. Interactome separation (S_AB) under harmonisation

| Disease pair | Reported | Harmonised N = 1,000 | Harmonised N = 2,720 |
|---|---|---|---|
| PS-AD | -0.267 | **-0.573** | **-0.538** |
| PS-HS | **-0.594** | -0.356 | -0.519 |
| AD-HS | -0.152 | -0.260 | -0.401 |
| AD-CSU | -0.132 | -0.202 | -0.380 |
| PS-CSU | 0.020 | -0.175 | -0.361 |
| VL-CSU | -0.018 | -0.120 | -0.350 |
| AD-VL | -0.039 | -0.084 | -0.318 |
| PS-VL | -0.272 | -0.131 | -0.313 |
| HS-CSU | 0.050 | -0.130 | -0.304 |
| HS-VL | -0.185 | -0.010 | -0.273 |

All ten pairings among PS, AD, HS, VL and CSU are negative under harmonisation at both set sizes, including PS-CSU and HS-CSU, which are marginally positive in the primary analysis. The finding that these five diseases occupy closer regions of the interactome than random expectation is therefore independent of the thresholds used, and CSU's marginal separation in the primary analysis reflects the smaller size of its module. As with gene overlap, PS-AD and PS-HS are the two closest pairs, separated by 0.019 at the largest set size.

#### 4. SLS-ICD

Three observations were reported as evidence that SLS-ICD is distinct: it shares none of the 14 genes common to the other five diseases, it shares few differentially expressed genes with them, and its S_AB is positive against every other disease. We have examined each. They are not independent, and two are governed by the size of the SLS-ICD gene set (33 genes).

Gene overlap, corrected for set size (hypergeometric expectation, 10,340 genes measurable on all platforms):

| Pair | Observed overlap | Expected by chance | Observed / expected |
|---|---|---|---|
| SLS-ICD-AD | 6 | 1.9 | **3.08** |
| SLS-ICD-CSU | 4 | 1.5 | **2.74** |
| SLS-ICD-PS | 19 | 8.6 | **2.20** |
| SLS-ICD-HS | 14 | 7.6 | **1.85** |
| SLS-ICD-VL | 3 | 5.7 | 0.53 |
| PS-HS (for comparison) | 1,993 | 1,397 | 1.43 |

Relative to its size, SLS-ICD shares more genes with PS, HS, AD and CSU than expected by chance, and proportionally more than PS shares with HS. The small absolute counts follow from SLS-ICD contributing 33 genes rather than several thousand. Similarly, the absence of the 14 shared genes from the SLS-ICD set is the expected outcome: drawing 33 genes from approximately 20,000, the expected number of those 14 recovered is 0.02. Neither observation supports SLS-ICD being unusually disconnected.

S_AB is likewise size-dependent. The reported positive values derive from a 33-gene module, whose internal distances are large because the genes are sparsely distributed across the interactome. When SLS-ICD is given an equally sized set for comparison with the other diseases, the values become negative (PS-SLS-ICD -0.129 at N = 582). Neither figure supports an inference about the position of SLS-ICD in the interactome, and we do not present one.

What does hold is the pathway evidence, which is unaffected by these considerations because enrichment testing accounts for the size of the query set. The 33 SLS-ICD genes form a coherent programme of cornified-envelope and keratinocyte-differentiation genes (KRT6B, KRT77, LCE5A, SPRR1A, SPRR2B, TGM3, S100A7, S100A7A, S100A8, IL36G), yielding 52 significant biological-process terms led by keratinocyte differentiation (p.adj = 2.9 x 10-7, 8 of 33 genes). The other five diseases are dominated by immune activation. The distinction between SLS-ICD and the clinical diseases is therefore qualitative, residing in which biological programme is engaged, and it rests on enrichment rather than on gene counts or module topology.

**Overlap under harmonization.** Under the harmonized procedure, in which every condition contributes an identically sized gene set, SLS-ICD continued to show the lowest overlap with the other conditions. Its mean pairwise overlap was 0.47 times that of the pairs not involving SLS-ICD, and this ratio was identical at all three set sizes examined. SLS-ICD pairs occupied the lowest ranks throughout, with VL-SLS-ICD the smallest overlap at every set size. This result is not attributable to gene-set size, which the harmonization removes.

| Pair | N = 250 | N = 400 | N = 582 | Rank at N = 582 (of |
|---|---|---|---|---|

| | | | | 15) |
|---|---|---|---|---|
| PS-SLS-ICD | 18 | 35 | 49 | 8 |
| HS-SLS-ICD | 13 | 26 | 41 | 11 |
| AD-SLS-ICD | 10 | 19 | 35 | 12 |
| CSU-SLS-ICD | 5 | 15 | 26 | 14 |
| VL-SLS-ICD | 4 | 9 | 24 | 15 |
| **Mean, SLS-ICD pairs** | **10.0** | **20.8** | **35.0** | **-** |
| **Mean, all other pairs** | **21.3** | **44.0** | **74.3** | **-** |
| **Ratio** | **0.47** | **0.47** | **0.47** | **-** |

#### 5. Summary

| **Finding** | **Status under harmonisation** |
|---|---|
| All five clinical diseases are more closely connected in the interactome than random expectation | **Confirmed at both set sizes** |
| PS-HS and PS-AD are the two most closely related pairs | **Confirmed by overlap and by S_AB at every set size** |
| PS-HS is the single most closely related pair | Depends on analysis depth; PS-HS leads on the complete gene lists, PS-AD at the top of ranked lists |
| VL has a distinct ribosome-biogenesis and RNA-metabolism profile | **Confirmed** |
| SLS-ICD engages a distinct biological programme (barrier repair and cornification) rather than immune activation | **Confirmed by enrichment, which is independent of set size** |
| SLS-ICD shares unusually few genes with the other diseases | Not confirmed; overlap exceeds chance expectation once set size is accounted for |
| SLS-ICD is topologically separated in the interactome (S_AB > 0) | Not interpretable; the value is governed by gene-set size |

#### 6. Materials provided

Harmonised gene sets, pairwise overlap and Jaccard values, S_AB values for every pair and set size, and the scripts used to generate them are provided in the accompanying data folder. S_AB was computed with the published Menche et al. (2015) separation algorithm on the unmodified human interactome (13,460 proteins, 141,296 interactions).
