## Supplementary_Table_S1_drug_candidates for "A Multilayer Human Skin Diseasome Links Molecular Architecture, Clinical Co-occurrence, and Drug-Reversal Signatures"

**Supplementary Table S1.** Drug-repurposing candidates shared between each skin condition and its TriNetX-observed comorbidities. Candidate comorbidities were derived from DisGeNET disease enrichment of each condition's differentially expressed gene set (Benjamini-Hochberg FDR < 0.05; FDR < 0.25 for VL, for which no association reached FDR < 0.05) and examined for patient co-occurrence in TriNetX. For each comorbidity, its DisGeNET-curated gene set was queried against the LINCS L1000 reference using unsigned gene-set enrichment (Fisher's exact test, |z| > 2). Shared candidates are compounds appearing among both the top 100 compounds so identified and the top 100 reversal candidates derived from the skin condition's own transcriptomic signature. TriNetX co-occurrence is the number of patients within the index-condition cohort who also carried the candidate diagnosis, relative to the cohort total; counts are rounded by TriNetX and the smallest reportable non-zero value is ten. Rows are ordered by skin condition, then by number of shared candidates.

| Skin condition | TriNetX-observed comorbidity | DisGeNET FDR | TriNetX co-occurrence | % of cohort | Shared candidates (n) | Shared candidate compounds |
|---|---|---|---|---|---|---|
| AD | Liver carcinoma | 1.75e-22 | 20/7610 | 0.26 | 25 | palbociclib;devazepide;WZ-3146;WZ-4-145;ZK-164015;NNC-05-2090;naproxol;estradiol-benzoate;WZ-4002;TG-101348;NTNCB;BRD-K31706415;mibefradil;tamoxifen;wortmannin;T-98475;thioridazine;BRD-K85211514;amlodipine;SU-11652;BMS-536924;tyrphostin-A9;Y-134;GSK-1070916;etoposide |
| AD | Dermatitis, Allergic Contact | 0.00171 | 290/7610 | 3.81 | 6 | aminopurvalanol-a;nobiletin;TG-101348;BRD-K87125912;purvalanol-a;JAK3-inhibitor-I |
| AD | Influenza | 8.98e-12 | 520/7610 | 6.83 | 6 | rimcazole;NTNCB;CAM-9-027-3;wortmannin;terfenadine;T-98475 |
| AD | Chronic Obstructive Airway Disease | 0.033 | 1830/7610 | 24.05 | 3 | tyrphostin-A9;hydrocortisone-valerate;CVF-SUMO-14 |
| AD | Obesity | 0.00203 | 1510/7610 | 19.84 | 3 | tyrphostin-A9;CAM-9-027-3;JAK3-inhibitor-I |
| CSU | HIV Infections | 4.79e-11 | 10/510 | 1.96 | 11 | methotrexate;mycophenolic-acid;BRD-K84959002;CCT-018159;PLX-4720;BRD-K02562327;mycophenolate-mofetil;lovastatin;PP-110;BRD-A24021119;hinokitiol |
| CSU | Dermatitis, Allergic Contact | 1.79e-07 | 30/510 | 5.88 | 8 | BIIB-021;aminopurvalanol-a;AZD-7762;TG-101348;KU-0060648;BRD-K87125912;purvalanol-a;AS-605240 |
| CSU | Atherosclerosis | 1.62e-05 | 50/510 | 9.8 | 5 | BRD-K87125912;tamoxifen;bosutinib;BMS- |

| | | | | | | |
|---|---|---|---|---|---|---|
| | | | | | | 536924;CCT-018159 |
| CSU | Pneumonia | 0.000251 | 80/510 | 15.69 | 4 | tamoxifen;GSK-1070916;midostaurin;saracatinib |
| CSU | Infarction, Middle Cerebral Artery | 0.000251 | 20/510 | 3.92 | 3 | nilotinib;PU-H71;fostamatinib |
| CSU | Osteoporosis | 0.00137 | 50/510 | 9.8 | 2 | PI-828;PP-110 |
| HS | Dermatitis, Allergic Contact | 3.98e-05 | 60/3490 | 1.72 | 6 | anisomycin;TG-101348;BRD-K93367411;alprenolol;riluzole;AS-605240 |
| HS | Spontaneous abortion | 2.38e-06 | 80/3490 | 2.29 | 3 | fraxetin;homoharringtonine;AS-605240 |
| HS | Multiple Sclerosis | 0.00974 | 50/3490 | 1.43 | 2 | AS-605240;TG-101348 |
| HS | Pneumonia | 0.000643 | 410/3490 | 11.75 | 1 | SA-1447005 |
| HS | Rheumatoid Arthritis | 3.9e-08 | 120/3490 | 3.44 | 1 | aminomethyltransferase |
| PS | Endometriosis | 0.0143 | 110/10140 | 1.08 | 7 | BMS-536924;AZD-8055;BRD-K62169556;midostaurin;vemurafenib;wortmannin;NVP-TAE684 |
| PS | Myocardial Ischemia | 0.000863 | 1430/10140 | 14.1 | 5 | ruxolitinib;T-98475;NU-7441;MK-2206;TPCA-1 |
| PS | Spontaneous abortion | 0.0143 | 40/10140 | 0.39 | 5 | BMS-536924;KU-0060648;KU-0063794;AS-605240;CCCP |
| PS | Influenza | 2.8e-07 | 520/10140 | 5.13 | 4 | BRD-K74623475;sirolimus;wortmannin;T-98475 |
| VL | Amyotrophic Lateral Sclerosis | 0.19 | 10/1180 | 0.85 | 3 | MPEP;trifluridine;olomoucine |
| VL | Optic Atrophy | 0.177 | 10/1180 | 0.85 | 3 | WT-171;BRD-K22052171;BRD-K05978741 |
| VL | Anemia, Diamond-Blackfan | 0.136 | 10/1180 | 0.85 | 2 | olomoucine;oxyphenonium |
| VL | Frontotemporal Lobar Degeneration | 0.136 | 40/1180 | 3.39 | 1 | WT-171 |

*Source data: signaturesearch/TableS_trinetx_corroborated_drug_candidates.csv*
