## Supplementary_Table_S2_datasets for "A Multilayer Human Skin Diseasome Links Molecular Architecture, Clinical Co-occurrence, and Drug-Reversal Signatures"

**Supplementary Table S2. Transcriptomic datasets analysed.**

All six datasets were obtained from the NCBI Gene Expression Omnibus and analysed with limma via GEO2R. Sample counts and group composition are those recorded in the GEO series records. A uniform effect-size criterion of |log2 fold change| ≥ 0.585 (1.5-fold) was applied throughout; the significance criterion differs by dataset because no atopic dermatitis or SLS-ICD gene satisfies both the fold-change criterion and an FDR-adjusted P < 0.05, as set out in Supplementary Data S1. "Features in the expression table" is the number of rows in the differential-expression output analysed for that dataset. Up- and downregulated counts are the unique gene symbols with positive and negative log2 fold change respectively. Gene symbols in the resulting lists reflect the platform annotation available at the time of analysis. For GSE168735, which contains several chemical exposures, the comparison analysed was SLS-exposed versus untreated control skin; this is confirmed by the archived differential-expression output, whose moderated-t total degrees of freedom (19.69) are consistent with 11 versus 8 samples and incompatible with the 36-sample vehicle group.

| **Condition** | **GEO accession** | **Platform** | **Samples in series** | **Sample groups in series** | **Contrast analysed** | **Significance criterion** | **Features in the expression table** | **DEGs (unique symbols)** | **Upregulated** | **Downregulated** |
|---|---|---|---|---|---|---|---|---|---|---|
| Psoriasis (PS) | GSE30999 | GPL570, Affymetrix HG-U133 Plus 2.0 | 170 | 85 lesional, 85 non-lesional | Lesional vs non-lesional (85 vs 85) | Nominal P < 0.05 | 54,675 | 6,522 | 3,499 | 3,023 |
| Atopic dermatitis (AD) | GSE32924 | GPL570, Affymetrix HG-U133 Plus 2.0 | 33 | 13 lesional, 12 non-lesional, 8 normal | Lesional vs non-lesional (13 vs 12) | Nominal P < 0.05 | 54,675 | 1,527 | 731 | 796 |
| Hidradenitis suppurativa (HS) | GSE148027 | GPL570, Affymetrix HG-U133 Plus 2.0 | 33 | 18 lesional, 7 non-lesional, 8 healthy | Lesional vs non-lesional (18 vs 7) | FDR-adjusted P < 0.05 | 54,675 | 6,801 | 3,509 | 3,292 |
| Vitiligo (VL) | GSE75819 | GPL6884, Illumina HumanWG-6 v3.0 | 30 | 15 lesional (depigmented), 15 non-lesional | Lesional vs non-lesional (15 vs 15) | FDR-adjusted P < 0.05 | 20,099 | 3,040 | 1,749 | 1,291 |
| Chronic spontaneous urticaria (CSU) | GSE57178 | GPL6244, Affymetrix Human Gene 1.0 ST | 18 | 6 patient lesional, 7 patient non-lesional, 5 normal | Lesional vs non-lesional (6 vs 7) | Nominal P < 0.05 | 33,297 | 976 | 804 | 172 |
| SLS-induced irritant contact dermatitis (SLS-ICD) | GSE168735 | GPL29838 | 103 | 11 SLS-exposed, 8 untreated, 36 petrolatum vehicle, remainder other chemical exposures | SLS-exposed vs untreated control (11 vs 8) | Nominal P < 0.05 | 21,201 | 33 | 16 | 17 |
